# Cochlear organoids reveal epigenetic and transcriptional programs of postnatal hair cell differentiation from supporting cells

**DOI:** 10.1101/2021.09.19.460948

**Authors:** Gurmannat Kalra, Danielle Lenz, Dunia Abdul-Aziz, Craig Hanna, Brian R. Herb, Carlo Colantuoni, Beatrice Milon, Madhurima Saxena, Amol C. Shetty, Ronna P. Hertzano, Ramesh A. Shivdasani, Seth A. Ament, Albert S. B. Edge

**Author notes:** Correspondence (AE). Co-first author (Gurmannat Kalra, Danielle Lenz).

## Abstract

We explored the transcriptional and epigenetic programs underlying the differentiation of hair cells from postnatal progenitor cells in cochlear organoids. Heterogeneity in the cells including cells with the transcriptional signatures of mature hair cells allowed a full picture of possible cell fates. Construction of trajectories identified *Lgr5+* cells as progenitors for hair cells and the genomic data revealed gene regulatory networks leading to hair cells. We validated these networks, demonstrating dynamic changes both in expression and predicted binding sites of these transcription factors during organoid differentiation. We identified known regulators of hair cell development, *Atoh1, Pou4f3*, and *Gfi1*, and predicted novel regulatory factors, *Tcf4*, an E-protein and heterodimerization partner of Atoh1, and *Ddit3*, a CCAAT/enhancer-binding protein (C/EBP) that represses *Hes1* and activates transcription of *Wnt* signaling-related genes. Deciphering the signals for hair cell regeneration from mammalian cochlear supporting cells reveals candidates for HC regeneration which is limited in the adult.

## INTRODUCTION

The mouse cochlea contains approximately 3,000 hair cells. Its dimensions and location, and the small number of hair cells, make mechanistic, developmental, and cellular replacement studies difficult. We recently published a protocol to expand and differentiate murine neonatal cochlear progenitor cells into 3-D organoids that recapitulate developmental pathways and can generate large numbers of hair cells with intact stereociliary bundles, mechanotransduction channel activity, and molecular markers of native cells, including markers for both inner and outer hair cells (1). We also showed that inner and outer hair cells were segregated into separate organoids based on hair cell markers, prestin and vGlut3 (1), suggesting that some of the phenotypic complexity of cochlear hair cell types was represented in this *in vitro* system.

The relevance of cochlear organoids to *in vivo* differentiation depends on the fidelity with which they mimic *in vivo* cell types and regenerative processes (2, 3). The organoids held promise for modeling development and differentiation and for higher-throughput screening to identify small molecules and genes that modulate these processes (4). However, the analyses were limited to a small number of known marker genes, which may not fully reveal the cells’ transcriptional states. Moreover, the transcriptional signatures of cells within organoids that do not become hair cells remain little characterized. Here, we performed a comprehensive transcriptional and epigenomic characterization of the cochlear organoids in comparison to *in vivo* cell types, confirming that the organoids mimic nearly all supporting cell and hair cell subtypes of the *in vivo* cochlea as well as the vestibular system. In addition, gene regulatory network modeling of these data predicts novel regulators of hair cell development and differentiation.

## RESULTS

### Data integration identifies marker genes for cochlear and utricular cell types to allow robust analysis of hair cell differentiation in cochlear organoids

We have recently developed organoids made by expansion of *Lgr5+* supporting cells from cochlear sensory epithelium (1). Since the conditions for expansion resulted in organoids that were 78% *Lgr5+*, and the differentiation steps resulted in a mix of cells that included inner and outer cochlear hair cells, we chose to analyze the phenotypic complexity represented by the organoid system to determine the epigenetic and transcriptional changes underlying the postnatal differentiation of hair cells from supporting cells.

To enable a quantitative comparison of inner ear sensory epithelial cell types, we developed a comprehensive database of cell type-specific gene expression in the mouse cochlea and vestibular organs (Fig. 1A). We generated new scRNA-seq from mouse utricle at postnatal days 2 and 7 (Fig. 1B and C). We integrated these data with six previously published scRNA-seq datasets to identify robust marker genes for 17 cell types, including subtypes of hair cells, supporting cells, spiral ganglion neurons, and cells of the stria vascularis (Supplemental Tables 1 and 2). We also combined all hair cells, supporting cells, spiral ganglion neurons and strial cells to determine shared molecular signatures among the subtypes of each of those categories. We correlated these markers with individual *in vivo* cell types over multiple developmental time points (E14-P7) as validation of the shared and specific molecular signatures between cell types (Fig. 1B, Supplemental Table 3). A plot of the expression of the top genes of each cell type revealed a robust correlation of genes within each cell type and state as compared to related cell types and states for hair cells, neurons, supporting cells and strial cells (Fig. 1C) and revealed which genes were expressed across related cell types.

**Figure 1.**
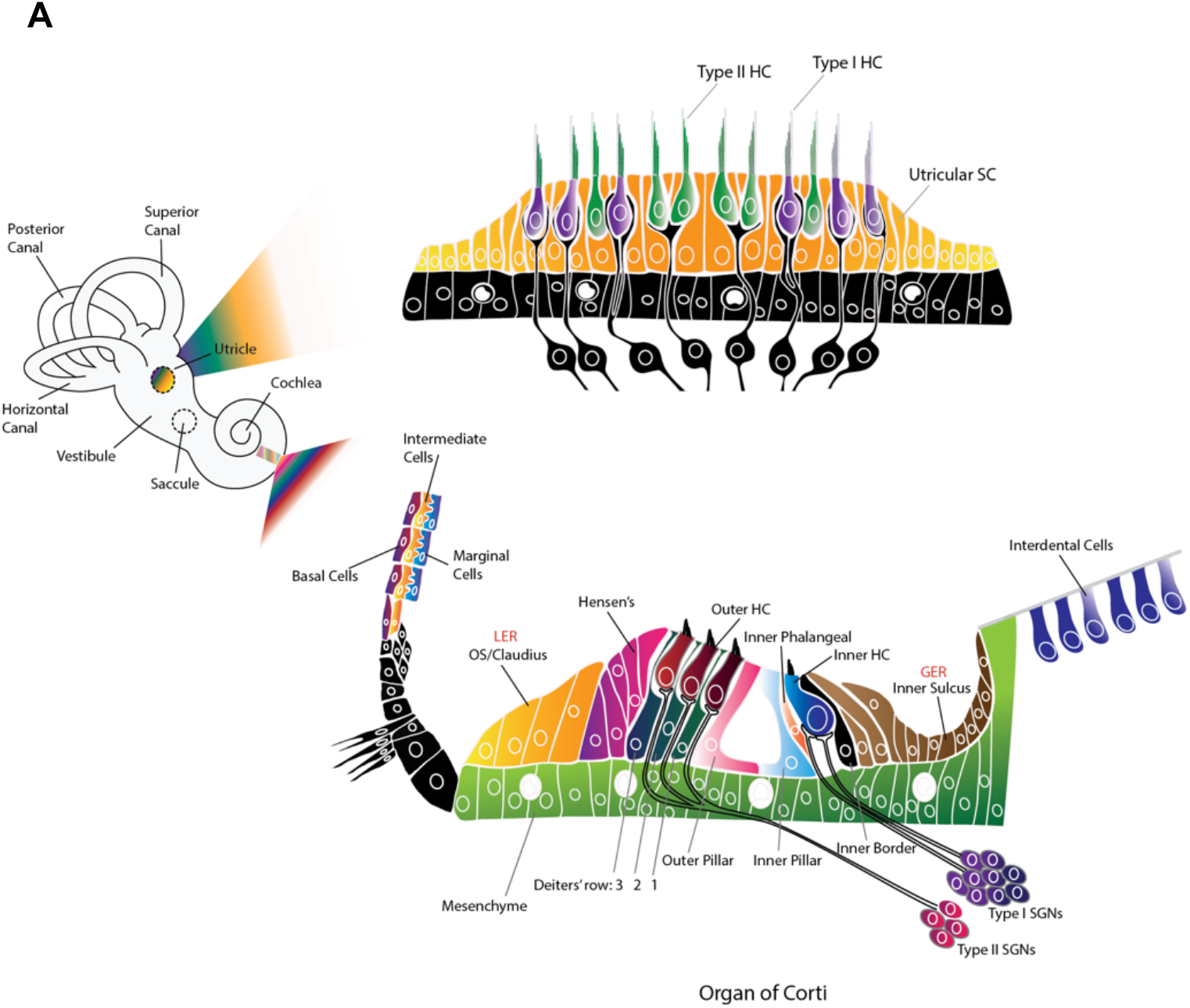
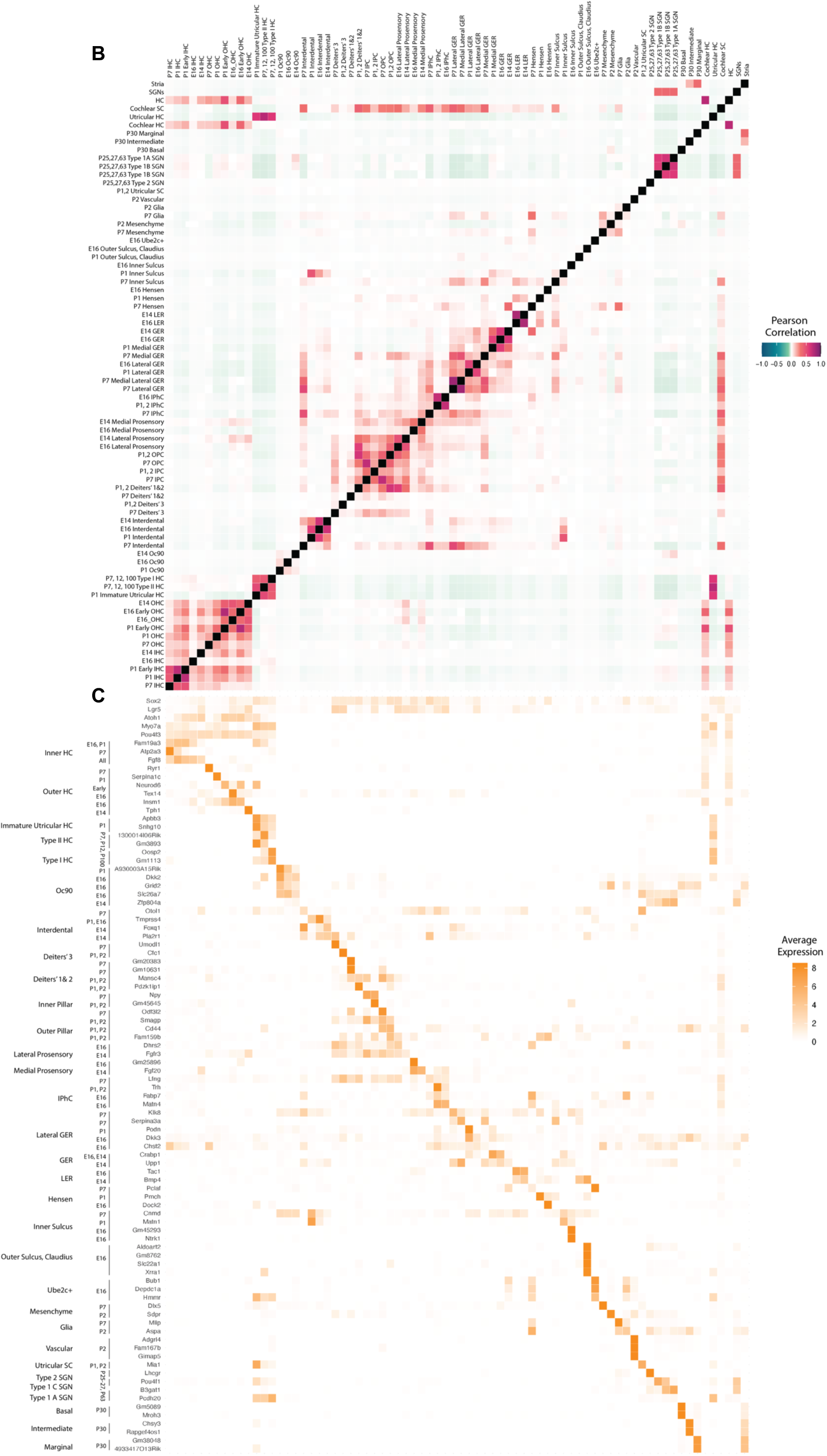
Marker genes for cochlear and utricular cell types derived by scRNA-seq integration. A. Anatomical organization of transcriptionally-defined cell types in the mammalian cochlea and utricle. Robust marker genes for each cochlear and utricular cell type were defined by integration of multiple scRNA-seq datasets and compared to marker genes identified at specific developmental timepoints (5). B. Pairwise correlations of marker genes across cell types based on Pearson correlations of cell type specificity scores for up to 300 genes per cell type. C. Expression patterns of individual marker genes in each cell type. Abbreviations used are: HC: Hair cell; SC: Supporting cell; IPhC: Inner phalangeal cell; IPC: Inner pillar cell; OPC: Outer pillar cell; GER: Greater epithelial ridge; LER: Lesser epithelial ridge; OS: Outer sulcus; SGN: Spiral ganglion neuron.

Notably, known markers for cochlear hair cells – many of which were derived from analyses of protein abundance – are not always suitable for these analyses, either because their transcript levels are less cell type-specific or because they are also expressed in vestibular hair cells. For instance, a well-known marker of cochlear IHCs, *Slc17a8*, was also highly expressed in type I and type II utricular hair cells. We concluded that our array of subtype and cell state gene profiles would allow us to identify *in vitro* cell types with a high degree of specificity.

### Single-cell analysis of organoids reveals tight correlation to the postnatal cochlea

To assess gene expression and chromatin accessibility during the differentiation of *Lgr5+* cells to hair cells, we sought to obtain epigenetic and transcriptional signatures of the cells. We analyzed the trajectories of individual cells in the organoids at the single cell level at various times of differentiation by scRNA-seq starting from the expanded inner ear progenitors (referred to here as D0 cells) and differentiating them for 10 days *in vitro* (referred to here as D10 cells). We compared the profiles of the differentiating postnatal supporting cells to the complex mosaic of sensory cells in the sensory epithelium. We generated scRNA-seq of a total of 67,162 cells from cochlear organoids at day 0 (4 samples) and day 10 (2 samples) of differentiation (Fig. 2A). Examination of canonical marker genes indicated that at both time points organoids were primarily composed of epithelial cells that expressed markers of hair cells (*Atoh1, Pou4f3, Myo7a*, and *Pvalb*) or supporting cells (*Sox2* and *Lgr5*) (Fig. 2B). Louvain clustering revealed 11 transcriptionally distinct cell types (Fig. 2C). As expected, several cell clusters were differentially abundant in D0 vs. D10 samples (Fig 2D, *top*), indicating that differentiation resulted in substantial changes in cell composition. We correlated signatures of *in vivo* cochlear and utricular cell types and states at various time points (5) to signatures of the 11 clusters of cells from the cochlear organoids (Fig. 2D, *middle*; Supplemental Table 4). As validation, the cell types correlated well with the individual *in vivo* cell types and time points (Fig. 2D, *middle*; Supplemental Table 4). Our molecular profiles also held up strongly in the *in vivo* data (Fig. 2D).

**Figure 2.**
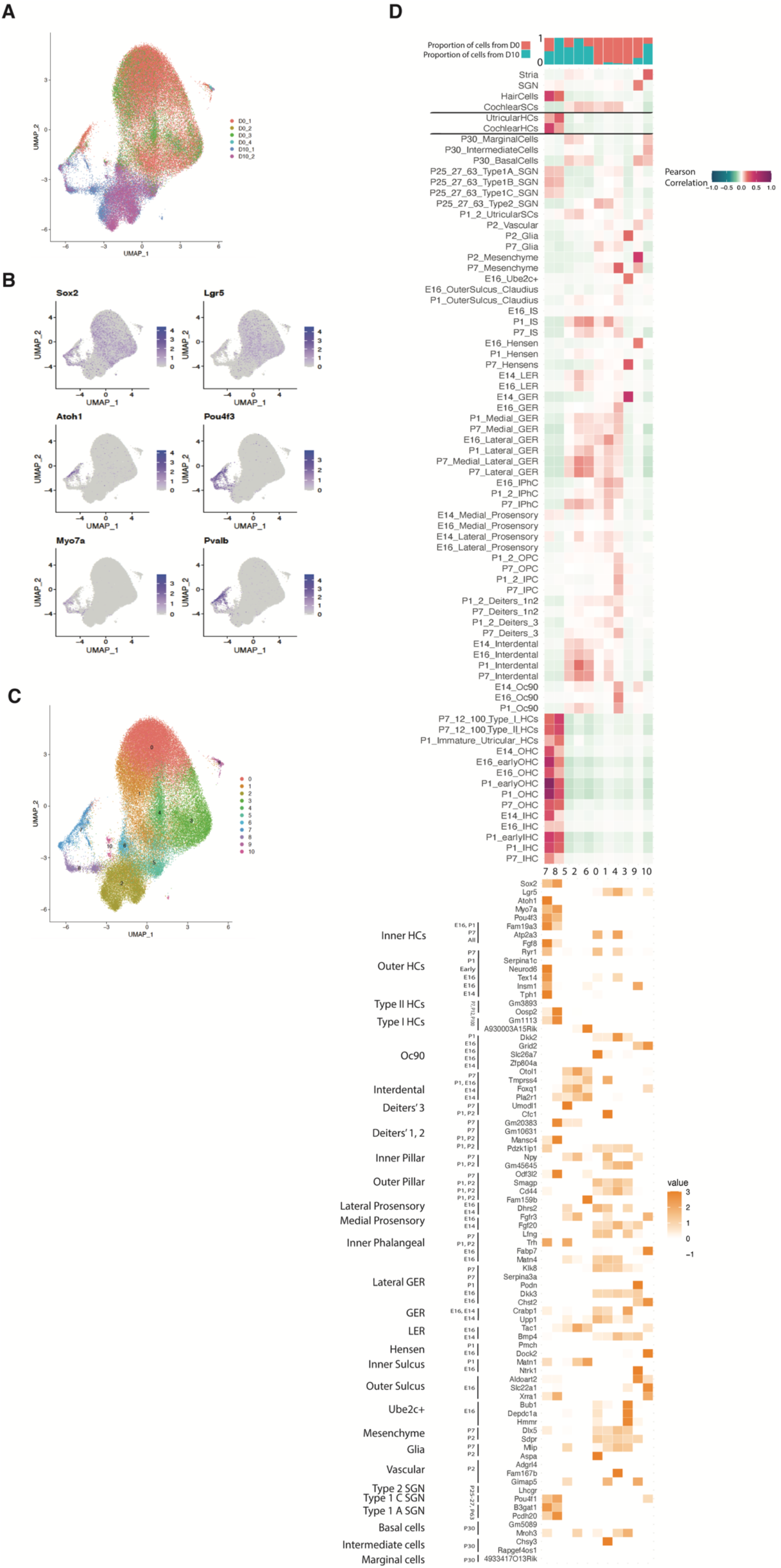
scRNA-seq characterization of *in vitro* cell types in cochlear organoids. A. UMAP of integrated scRNA-seq data from days 0 and 10 of organoid differentiation, labeled by sample and time point B. Expression of the canonical hair cell and supporting cell markers *Sox2, Lgr5, Atoh1, Pou4f3, Myo7a*, and *Pvalb*. C. UMAP labeled by Seurat clusters. D. *Top*: Proportion of cells in each in vitro cluster from day 0 vs. 10 of differentiation. *Middle*: Pairwise correlations of marker genes of *in vitro* clusters vs. *in vivo* cochlear and utricular cell types. Color intensity indicates Pearson correlations of cell type specificity scores for up to 300 genes per cell type. *Bottom*: Expression of top marker genes from *in vivo* cell types in *in vitro* clusters. For an interactive version of the trajectory analysis see https://umgear.org/Lgr5org.

Among the organoid clusters that lack hair cell markers, clusters 0, 1, 3, and 4 (primarily D0 cells) were positively correlated with signatures for *in vivo* inner phalangeal cells, inner pillar, GER, prosensory, inner sulcus, and *Ube2c*+ cells. The signature for cluster 9 (primarily D0 cells) was strongly correlated with Oc90 and mesenchymal cells. Clusters 2, 5, and 6 (primarily D10 cells) were positively correlated with signatures for postnatal lateral GER, interdental cells, and *Oc90*+ cells. Cluster 10 was correlated with the signature for Deiters’, *Oc90*+, inner pillar, and cells of the stria vascularis. Importantly, most of the *in vivo* cell types matching clusters from D0 cells expressed *Lgr5* – in particular, inner pillar, inner phalangeal cells, and lateral GER – and likely reflect the original cell types from which the organoids were derived. By contrast, none of the *in vivo* cell types matching clusters from D10 cells expressed *Lgr5*, suggesting that by day 10 of differentiation these cells have assumed new cell identities. These results indicate that cells within organoids represent nearly all supporting cell subtypes from the *in vivo* cochlea.

Signatures of two clusters – 7 and 8 – correlated strongly with *in vivo* hair cells. These cells represent 1.6% of the cells in D0 organoids and 12.4% of D10 organoids. Cluster 7 (49.9% D0 and 50.1% D10 cells) correlated most strongly with signatures of early cochlear hair cells. By contrast, cluster 8 (99.7% D10 cells) correlated less strongly with early cochlear hair cells, but more strongly with mature hair cells. We speculated that the small number of hair cells in D0 organoids arose through activation of hair cell differentiation programs, whereas more numerous and more mature hair cells arose during the 10-day course of organoid differentiation.

We examined multi-gene signatures and specific marker genes for subtypes of organoid-derived hair cells (Fig. 2D, *bottom*). Subsets of hair cells from the organoids showed genes differentially expressed in inner and outer hair cells, consistent with our previous report. Surprisingly, some organoid-derived hair cells also expressed signatures of type I and type II utricular hair cells, suggesting that these cells have not assumed a distinct cochlear vs vestibular identity (6). In addition, the canonical marker of immature hair cells, *Atoh1*, was expressed most strongly in cluster 7, while markers of more mature hair cells, including *Pou4f3* and *Myo7a*, were most highly expressed in cluster 8. Therefore, these clusters may correspond to immature and mature hair cell identities, respectively. Many of the organoid-derived hair cells expressed the progenitor cell marker *Sox2*. With the exception of utricular type II hair cells, *Sox2* is generally absent from hair cells; however, *Sox2* is expressed in newly generated hair cells, both in the embryo (7) and in regenerated hair cells that arise *in vivo* after noise- or chemically induced hair cell ablation (7, 8). In summary, differentiation results in a trajectory from immature to more mature hair cells of which subsets express markers for both cochlear and vestibular subtypes.

### Trajectories of supporting cell to hair cell transdifferentiation

Projecting the average expression of differentially expressed genes from hair cells at E14, E16, P1, and P7 [45], and from vestibular immature and mature hair cells revealed a hair cell maturation trajectory from organoid clusters 7 to 8 (Fig 3A). To further investigate the molecular differences between clusters 7 and 8, a differential gene expression analysis revealed the upregulation of cochlear and vestibular hair cell genes in cluster 8, and mostly cochlear hair cell genes in cluster 7 (Fig 3B). None of the hair cells expressed genes of other *Atoh1*-dependent lineages such as cerebellar granular precursors, intestinal epithelial cells, or Merkel cells (Fig. 3C).

**Figure 3.**
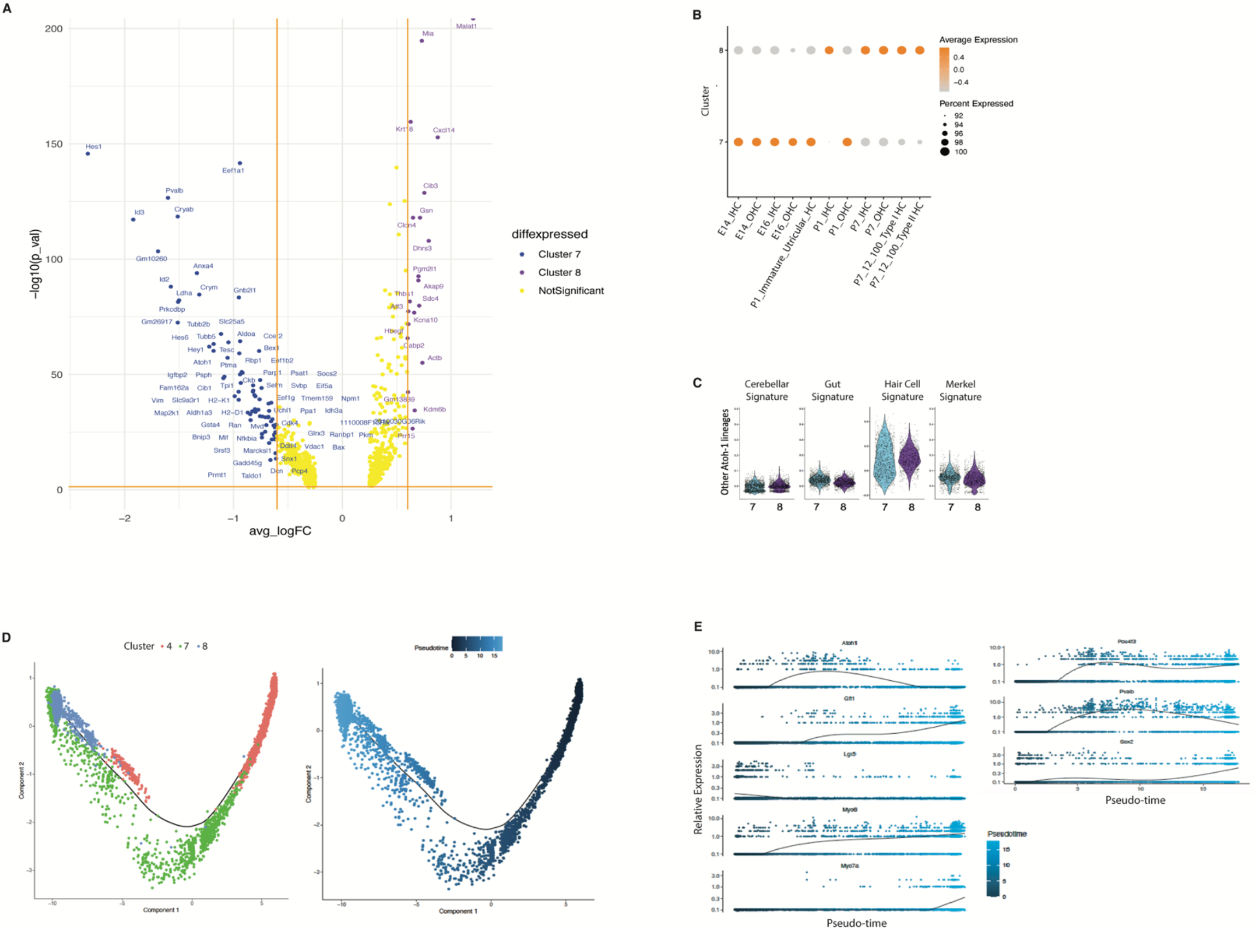
Molecular characterization and pseudotemporal trajectories of hair cell-like cells in cochlear organoids. A. Volcano plot indicating genes differentially expressed between two clusters of hair cell-like cells in organoids: cluster 7 (primarily in day 0 organoids) and cluster 8 (primarily in day 10 organoids). B. Aggregate expression of markers specifically expressed in E14, E16, P1, or P7 cochlear hair cells in clusters 7 and 8. Yellow indicates high expression, whereas dark red indicates low expression. C. Aggregate expression of genes expressed in cochlear and non-cochlear *Atoh1-*dependent cell lineages in clusters 7 and 8. D. Monocle pseudotime trajectory delineates transdifferentiation of *Lgr5+* supporting cells (cluster 4) to hair cell-like cells (clusters 7 and and 8). Trajectory labeled by cluster number (*left*) or pseudotime (*right*). For an interactive version of the trajectory analysis see https://umgear.org/Lgr5org. E. Expression of known marker genes with dynamic expression across pseudotime.

Next, we sought to model the trajectories by which *Lgr5+* supporting cells transdifferentiate to hair cells, using pseudotime trajectory analysis of our scRNA-seq data, as well as bulk mRNA sequencing from organoids at days 0, 2, 4, and 10 of differentiation. Using Monocle, we produced a non-branching trajectory from differentially expressed genes (Supplemental Table 5) in clusters 4, 7, and 8 that followed cells from cluster 4 (the cluster with the highest expression of Lgr5) through cluster 7 (immature hair cells) to cluster 8 (mature hair cells) (Fig. 3D).

Examination of known markers for hair cell development revealed sequential patterns of activation. Lgr5 expression was high at the beginning of the trajectory, then declined to low levels in immature and mature hair cells (Fig. 3E). Atoh1, the master regulator of hair cell development, peaked in immature hair cells at the middle of the trajectory. Markers of mature hair cells, including *Gfi1, Myo6, Myo7a, Pou4f3*, and *Pvalb* increased to high levels at the end of the trajectory. As noted above, Sox2 was co-expressed with mature hair cell markers in the organoids; its expression was highest at the end of the pseudotime trajectory. Extending this analysis to additional genes with dynamic expression across pseudotime, we identified 6,523 dynamically expressed genes. Clustering these genes with BEAM revealed three gene co-expression clusters (Supplemental Table 6).

To confirm these gene dynamics, we generated mRNA-seq of bulk organoid cells at days 0, 2, 4, and 10 of differentiation (Fig. 4A), as well as of P2 hair cells (*Atoh1*+) and supporting cells (*Lgr5*+ and *Sox2*+) (Fig. 4B). Principal component analysis on the D0, 2, 4, and 10 samples confirmed that the largest component of variation separating the cells were the different time points (Fig. 4A). Examination of known markers confirmed activation of hair cell marker genes during differentiation (e.g., *Myo7a* and *Tmc1*), accompanied by a decrease in Notch pathway genes (e,g., *Notch3* and *Jag1) and* cell cycle genes (e.g., *Ccnc1, Birc5*, and *Cdk1*; Fig. 4C). K-means clustering of the bulk RNA-seq data revealed eight distinct expression patterns (Fig. 4D, Supplemental Table 7). Three of the eight bulk RNA-seq -derived patterns statistically overlapped the three transdifferentiation-related co-expression patterns derived from the scRNA-seq trajectory (groups 1, 3, and 4) (Supplemental Table 8). Representative genes from these clusters include *Ccnd1* and *Lgr5* (group 1, decreasing expression during differentiation), *Atoh1* and *Pax2* (group 4, middle-onset expression), and *Myo7a* and *vGlut3* (group 3, late-onset expression) (Fig. 4E).

**Figure 4.**
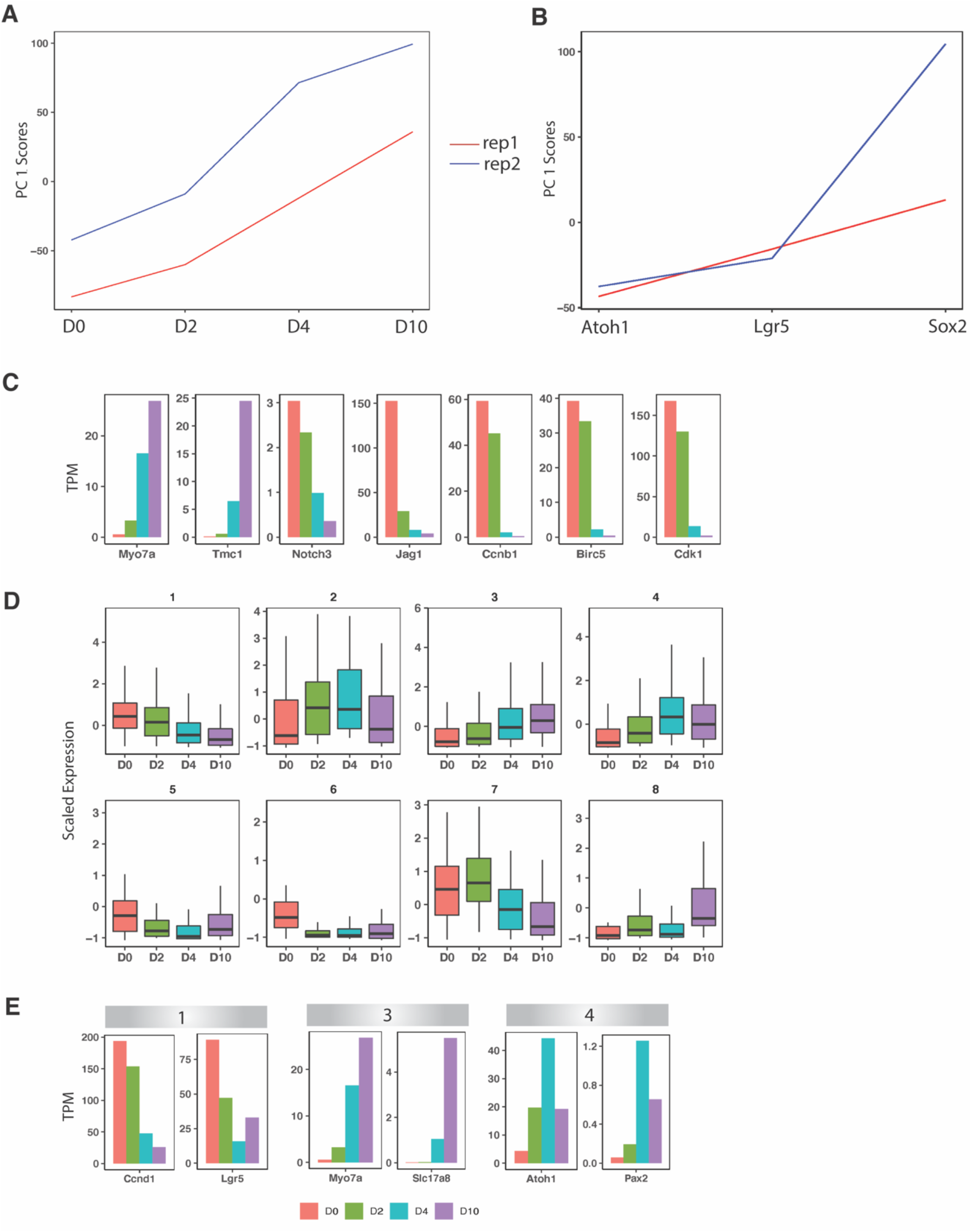
Clustering analysis of bulk RNA-seq data from D0, 2, 4, and 10. A. PCA of the D0, 2, 4, and 10 bulk RNA-seq samples and their replicates. B. PCA of *Atoh1*+, *Lgr5*+, and *Sox2*+ cells and their replicates from P2 mouse cochlea. C. Expression of select genes in D0, 2, 4,and 10 samples. D. K-means clustering of D0, 2, 4, and 10 bulk RNA-seq samples, showing eight gene expression patterns. E. Expression of select genes in groups 1, 3, and 4 from D.

Functional annotation of gene co-expression modules across datasets revealed biological processes that were robustly enriched at the early-, middle-, and late phases of transdifferentiation. Significant GO terms (p.adj < 0.05) for the “early” gene co-expression modules include negative regulation of Notch signaling pathway, regulation of inner ear auditory receptor cell differentiation, and regulation of mechanoreceptor differentiation. Significant GO terms for middle-onset module included “regulation of epithelial cell differentiation” (p = 4.01 × 10^−5^) and development-related terms. Significant GO terms for the late-onset module included cilium movement, inner ear development, hair cell differentiation, and inner ear morphogenesis (Supplemental Table 9). These gene expression dynamics largely mirror hair cell development processes observed *in vivo*.

### Gene regulatory networks underlying transdifferentiation

The transcriptional regulators involved in transdifferentiation of supporting cells to hair cells remain incompletely characterized. We predicted these regulators by reconstructing a gene regulatory network (GRN) model from our scRNA-seq and mRNA-seq data. Briefly, we applied a random forest GRN reconstruction algorithm, GENIE3 (9), to predict target genes for transcription factors, using scRNA-seq data from clusters 4, 7, and 8. This resulted in a GRN model predicting the regulation of 17,849 genes by 1024 TFs. To predict key regulators of transdifferentiation, we tested for the enrichment of GENIE3-derived regulons in the three modules (early-, middle-, and late-onset gene co-expression; Fig. 4E, Supplemental Table 8) observed in the bulk RNA-seq-derived patterns that overlapped the transdifferentiation-related co-expression patterns derived from the scRNA-seq trajectory. A total of 67 TFs were reproducibly enriched (p < 0.05) in modules derived from both scRNA-seq and bulk RNA-seq data (Fig. 5A). Next, these 67 TFs were used as input to reconstruct a dynamical TF-to-TF gene regulatory network along pseudotime, using SCODE (10) (Supplemental Fig. 1, Fig. 5B). TFs with the most predicted targets in this network include known regulators of hair cell differentiation (e.g., *Atoh1, Pou4f3*, and *Lmo1*), as well as TFs that have not previously been implicated in this process (e.g., *Ddit3, Basp1, Tcf4, Sox4*).

**Figure 5.**
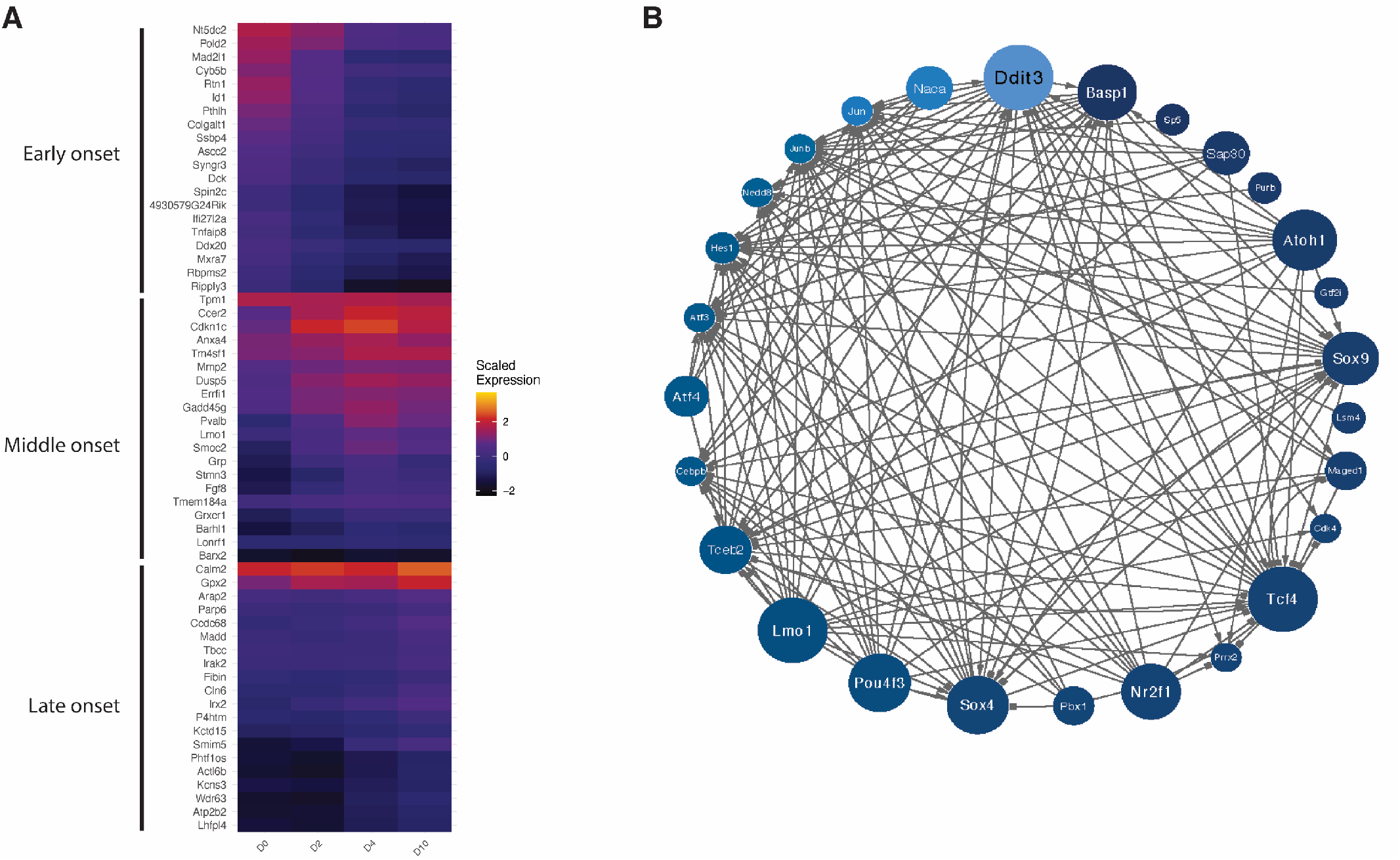
Gene regulatory network of transcription factors that drive the transdifferentiation of *Lgr5*+ cochlear progenitor cells to hair cells. A,B. Each transcription factor is colored by its activity (A) in the pseudotime course from Figure 3E, and its size in the diagram (B) reflects the magnitude of its outdegree.

### Chromatin accessibility changes associated with trans-differentiation of supporting cells to hair cell-like cells in cochlear organoids

We performed the Assay for Transposase-Accessible Chromatin (ATAC-seq) in cochlear organoids at days 0, 2, and 10 of differentiation to characterize changes in chromatin accessibility during the transdifferentiation of supporting cells to hair cells (n=2 biological replicates per condition). For comparison, we also generated ATAC-seq of sorted *Sox2+* and *Lgr5+* supporting cells and *Atoh1+* hair cells from *in vivo* mouse cochlea at postnatal day 1. Peak-calling with MACS identified 44,540 chromatin accessibility peaks that were reproducible across two or more samples (Supplemental Table 10). 15,123 of these peaks showed at least a nominally significant change in accessibility across groups (one-way ANOVA, *P* < 0.05). We used these data to explore gene regulatory mechanisms underlying transdifferentiation, focusing on TFs implicated in our gene expression-based gene regulatory network model.

First, we considered the patterns of chromatin accessibility at the promoters of predicted key regulator TFs. The promoter regions for many of these TFs were differentially accessible during transdifferentiation, with varying dynamics. For instance, the *Atoh1* promoter was already accessible at day 2, whereas the *Pou4f3* promoter became accessible primarily at day 10 (Fig. 6A,B). As expected, the promoters of *Atoh1* and *Pouf43* were also accessible in *Atoh1+* sorted hair cells from the *in vivo* cochlea, but not in *Lgr5+* or *Sox2+* supporting cells. By contrast, the promoter regions of several transcription factors predicted to regulate the early stages of transdifferentiation, including *Sox9* and *Atf3*, showed decreased accessibility from day 0 to day 10, as well as in *Atoh1+* hair cells vs. *Lgr5+* and *Sox2+* supporting cells (Fig. 6D,E). In addition, the promoters of some TFs with dynamic gene expression across transdifferentiation had similar chromatin accessibility throughout this process (e.g., *Tcf4*, Fig. 6C). These results suggest that transdifferentiation involves cis-acting changes in the chromatin states of genes encoding key regulator TFs.

**Figure 6.**
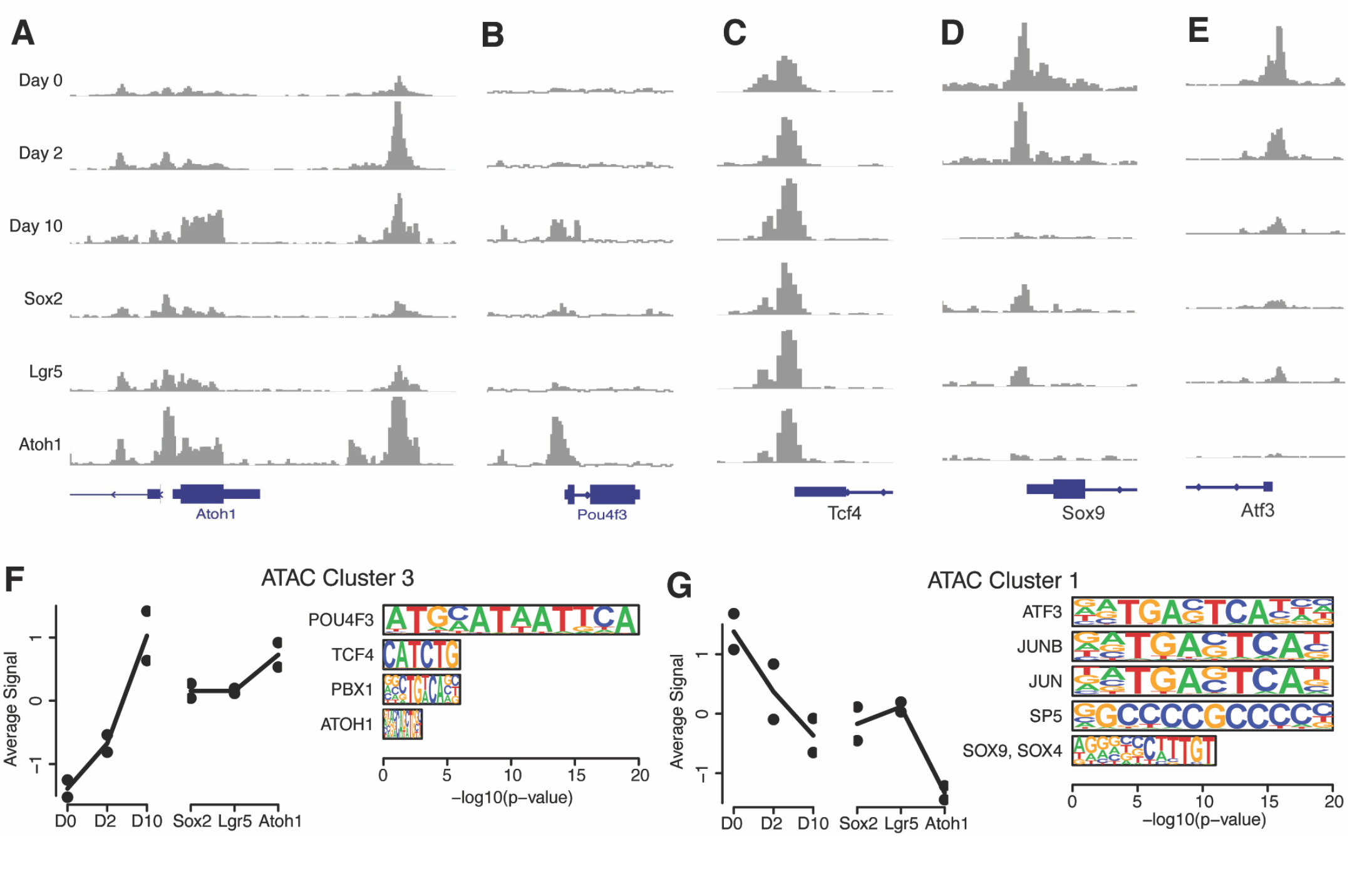
Chromatin accessibility dynamics during transdifferentiation in cochlear organoids. A-E. Chromatin accessibility at the promoters of the known and predicted key regulator transcription factors *Atoh1* (A), *Pou4f3* (B), *Tcf4* (C), *Sox9* (D), and *Atf3* (E). F-G) Average accessibility pattern for peaks within clusters 3 (F) and 1 (G), and enrichment of peaks for sequence motifs recognized by key regulator TFs with dynamic expression in the RNA-based gene regulatory network model.

Next, we characterized trans-acting effects of these TFs on chromatin accessibility. Using k-means clustering, we identified 12 patterns of chromatin co-accessibility across our *in vitro* and *in vivo* data (Supplemental Fig. 2; Supplemental Table 10). Several of these patterns describe increasing or decreasing chromatin accessibility across trans-differentiation *in vitro*, accompanied by concordant differences of chromatin accessibility in hair cells vs. supporting cells *in vivo*. For instance, peaks in Cluster 3 were up-regulated at day 10 of transdifferentiation, as well as in *Atoh1+* hair cells (Fig. 6F), while peaks in Cluster 1 were down-regulated at day 10, as well as in *Atoh1+* hair cells (Fig. 6G). Sequence motif enrichment analysis with HOMER (11) predicted TFs that may regulate these patterns (Supplemental Table 11). Importantly, clusters characterized by dynamic changes in chromatin accessibility across transdifferentiation were enriched for motifs recognized by several of the key regulator TFs from our gene regulatory network model, providing independent validation. Specifically, Cluster 3 (up-regulated in hair cells) was enriched for Octamer motifs recognized by Pou4f3, as well as E-box motifs recognized by Atoh1 and Tcf4 (Fig. 6F). Cluster 1 (down-regulated) was enriched for SRY-box motifs recognized by Sox family transcription factors, as well as motifs recognized by several activity-dependent factors such as Atf3, Jun, and Junb (Fig. 6G). Changes in chromatin accessibility governed by key regulator TFs with dynamic expression may therefore regulate the dynamic activity of thousands of enhancers and promoters in this context.

## DISCUSSION

A capacity for transdifferentiation of sensory epithelial supporting cells to hair cells allows the chick to regenerate hair cells in the deafened cochlea. Although cochlear sensory epithelial supporting cells in the adult mammal lack the capacity for regeneration, the cells show the capacity to differentiate into hair cells in the early postnatal period (7, 12, 13). Deciphering the signals for reprogramming of mammalian cochlear supporting cells to hair cells would be an important step toward therapies for hair cell regeneration as a treatment for deafness. Here, utilizing an established protocol (1), we generated cochlear organoids from murine *Lgr5+* progenitor cells and performed a comprehensive molecular characterization at multiple time points in their differentiation by scRNA-seq, bulk RNA-seq, and ATAC-seq. Transcriptional signatures of mature hair cells were apparent after ten days of organoid differentiation and found that during the course of differentiation the cells mimicked nearly all subtypes of supporting cells and hair cells in the newborn cochlea. From these data, we reconstructed a gene regulatory model to gain insight into the transcriptional and epigenetic programs that drive the differentiation of *Lgr5+* progenitor cells to a hair cell fate.

Clustering of the cells in the UMAP allowed us to identify several groups of cells that were derived largely from the day zero time point of organoid differentiation and were related to supporting cells and the surrounding epithelium in the cochlear duct. Tracing their lineage to the hair cell clusters showed that cluster 4, which expressed *Lgr5* and *Sox2*, was the primary source of hair cells and allowed us to identify genes in their trajectory from supporting cell to hair cell. The cells at the start of the differentiation protocol correlated best with *in vivo* inner and outer pillar, inner phalangeal, Deiters’, prosensory, greater epithelial ridge, and inner sulcus cells. Cells made by a variant of this protocol were largely comprised of GER cells (14), and organoids grown from human cochlear cells also expressed markers of both supporting cells and hair cells (3, 15). Cells at the end of the differentiation protocol correlated best with hair cells.

We validated these networks across transcriptional and epigenomic datasets. Dynamic changes in TF expression were confirmed across bulk and single-cell RNA sequencing datasets, and in many cases were accompanied by changes in the accessibility of each TF’s promoter. Trans-acting effects of these same TFs on downstream target genes were predicted from TF-gene co-expression, as well as by the enrichment of their sequence motifs in networks of co-accessible chromatin regions. We also integrated the *in vitro* data with six previous studies of cochlear and utricular cell types and with newly generated *in vivo* data from intact cochlea and utricle.

We present here a thorough database of novel, robust marker genes for cell types of the cochlea and the utricle, which allowed us to identify cell types involved in the *in vitro* organoid differentiation protocol. We also present gene expression dynamics during the organoid differentiation protocol, supported by scRNA-seq and bulk RNA-seq data. Our data integration analysis, which yielded robust cell type-specific marker genes for various cochlear and utricular cell types allowed a more precise characterization of cells involved in the organoid differentiation protocol. At later time points, hair cells from organoids expressed mixed transcriptional signatures for cochlear and vestibular subtypes. We show for the first time here that the expanded Lgr5+ cells in the cochlear organoids have the capacity to differentiate into cochlear and vestibular hair cell types with some cells assuming a mature hair cell identity. Further resolution of the progenitors and the resulting hair cells will be required to define the signaling that generates these diverse identities.

We previously demonstrated that organoid differentiation yielded cells expressing key hair cell markers, including those of inner and outer hair cells. Our analysis demonstrates that the hair cells reach maturity comparable to *in vivo* postnatal day 7. Our model identified known regulators of hair cell development, including *Atoh1, Pou4f3*, and *Gfi1*. It also predicted roles in postnatal hair cell differentiation for hair cell expressed genes, *Sox4, Tceb2, Nr2f1* and *Lmo1*. Their expression is consistent with previous findings on these genes. *Sox4* restores supporting cell proliferation and hair cell production after hair cell loss (16), *Tceb2* is expressed in cochlear hair cells (17), and *Nr2f1* (*COUP-TFI*) knockout has a significant increase in hair cell number and is thought to act through misregulation of Notch signaling components, including *Jag1, Hes5* and correlated with increases in supporting cell differentiation after inhibition of Notch activity in *COUP-TFI-/-* cochlear cultures (18). *Lmo1*: *Lim domain only 1*, is a transcriptional regulator that contains two cysteine-rich LIM domains but lacks a DNA-binding domain. *Lmo1*is specifically expressed in vestibular and cochlear hair cells (19).

Our exploration of the transcriptional network also elucidated the regulation of genes that had not been known to be expressed during supporting cell differentiation to hair cells. These include previously known patterns of hair cell and supporting cell genes as well as several new genes that follow those patterns. Upregulation of *Ddit3*, a cell cycle-related gene corresponding to endoplasmic reticulum stress, is a member of the CCAAT/enhancer-binding protein (C/EBP) family of transcription factors (20). It acts as a dominant-negative inhibitor by forming heterodimers with other C/EBP members, and as an inhibitor of the canonical Wnt signaling pathway by binding to Tcf7l2, impairing its DNA-binding properties and repressing its transcriptional activity (21). *Ddit3* loss, moreover, contributes to hearing loss (22). Our network reconstruction revealed that *Ddit3* had a high out-degree during differentiation, was most active towards the end of the maturation process, and acted similarly to *Pou4f3, Atoh1*, and *Lmo1* in repressing *Hes1* and activating transcription of Wnt signaling-related genes such as *Sox4, Tceb2, Jun* and *Junb* (20, 21). Another TF identified in our transcriptional network, *Atf4*, is a downstream target of *Ddit3*. It regulates expression of genes involved in endoplasmic reticulum function, reactive oxygen species production, and cell death (23). Among *Atf4* - target genes is C/EBP homologous protein, CHOP/GADD153 (24) which inhibits canonical Wnt signaling by interfering with binding of β-catenin to its interaction partners.

Our analysis also predicts novel regulatory factors such as *Tcf4*, the E-protein and heterodimerization partner of Atoh1. *Tcf4* interacts with Atoh1 to induce neural differentiation (25) but has not been reported in hair cell maturation literature, and was not considered essential as Atoh1 can interact with other E -proteins depending on cell context (25). Its high level of connection with other network genes indicates an important role in the control of hair cell differentiation. *Tcf4* mutations are causal for Pitt-Hopkins syndrome (26, 27), which is thought to be due to incomplete maturation or absence of cortical neurons. Tcf4 recruitment coincides with areas of high transcriptional activity as shown by the occurrence of H3K27Ac marks in regions adjacent to Tcf4 binding (26). This is a novel insight into a potentially fundamental role of a known transcription factor and interaction partner with Atoh1.

The progenitors are heterogeneous and consist of cells from GER as well as inner pillar and 3rd Deiters’ and other supporting and non-supporting cells that expand in the GSK3β inhibitor. The results show that *Lgr5*+ cells act as progenitors to hair cells. They are reprogrammed to hair cells by the combined activity of the γ-secretase inhibitor and GSK3β inhibitor to inhibit Notch and activate Wnt which stimulated the expression of several novel TFs and modeled *in vivo* postnatal progenitor cell differentiation to hair cells (7, 28-31). Their molecular trajectories adhered to the same overall steps as postnatal supporting cells and allowed a determination of epigenetic and transcriptional steps in their reprogramming to hair cells.

## METHODS

### Mice

Cells from the organ of Corti were prepared as described (1) from *Atoh1-nGFP* (32), *Sox2-GFP* (33), or *Lgr5-GFP* (34) mice. All animal procedures were approved by the Massachusetts Eye and Ear IACUC.

### Marker genes for each cell type

By integrating publicly available scRNA-seq data from cochlear and utricular cell types, we derived marker genes for each cell type. Utricular hair cells and cochlear hair cells and neurons were integrated into one expression matrix, and utricular supporting cells, cochlear supporting cells, and stria vascularis cells were integrated into a separate expression matrix. Marker genes were derived by performing differential gene expression analysis on each cell type against all others in that matrix, using the Seurat R package, and calculating a specificity score for each gene (5).

Datasets for hair cells and spiral ganglion neurons (SGNs):

| Cell Type | Time point | Source |
| --- | --- | --- |
| IHCs | P1, P7 | GSE137299 |
| OHCs | P1, P7 | GSE137299 |
| Type I HCs | P7, P12, P100 | GSE115934 |
| Type II HCs | P7, P12, P100 | GSE115934, GSE172327 |
| Type 1 SGNs | P63, P25-27 | GSE114759, GSE114997 |
| Type 2 SGNs | P63, P25-27 | GSE114759, GSE114997 |

Datasets for supporting cells and stria vascularis cells:

| Cell Type | Time Point | Source |
| --- | --- | --- |
| Hensen's | E16, P1 | GSE137299 |
| Inner Pillar | E16, P1, P2, P7 | GSE137299, GSE172327 |
| Outer Pillar | P1, P2, P7 | GSE137299, |
| Inner Phalangeal | E16, P1, P2, P7 | GSE137299, |
| Claudius | E16, P1 | GSE137299 |
| Deiters row 1,2 | P1, P2, P7 | GSE137299, |
| Deiters row 3 | P1, P2, P7 | GSE137299, |
| Utricular SCs | P1, P2 | GSE71982, GSE172327 |
| Marginal | P30 | GSE136196 |
| Intermediate | P30 | GSE136196 |
| Basal | P30 | GSE136196 |

Genes with a specificity score for the relevant cell type and either 0 for other cell types or a very low score for other cell types (as necessary for type 1A-C SGNs) were selected as marker genes.

### Expansion and differentiation of Lgr5+ cochlear progenitor cells

Inner ear organoids were made by expanding cochlear sensory epithelial cells by the established protocol (1) from newborn Atoh1-nGFP mice. The organ of Corti was dissected in medium supplemented with growth factors and expanded in GSK3β inhibitor, CHIR, and HDAC inhibitor, VPA, which results in the growth of *Lgr5+* cochlear hair cell progenitors. Organoids were induced to differentiate in GSK3β inhibitor, CHIR, and γ-secretase inhibitor, LY411575. Generation of hair cells was assayed by flow cytometry for GFP, which relies on Atoh1 enhancer activation in these transgenic mice (32).

### RNA-sequencing

Organoids were examined by RNA-Seq at 4 time points: the start (D0), the early (D2), middle (D4) and late (D10) stages of differentiation. RNA was isolated by published procedures (29, 35). The cells at these time points were compared to sorted cells prepared at P2 from *Atoh1-nGFP, Sox2-GFP* (33), and *Lgr5-GFP* (34) mice, corresponding to hair cells, supporting cells, and the *Lgr5+* subset of supporting cells. RNA quality was confirmed, and cDNA synthesis, and library preparation carried out using 2 ng of sheared cDNA. Illumina NextSeq500 Single-End 75 bp (SE75) sequencing was performed to provide an estimated coverage of 20-30 million single-end reads per sample as described (29).

### ATAC-sequencing

Concurrently we performed ATAC-Seq to query chromatin accessibility under these conditions. We performed these experiments using NextSeq500 Paired-End 40 bp (PE40) sequencing (Dana-Farber Cancer Institute Core). Initial data analysis, including alignment to the mouse genome (Mm10) was performed using Bowtie2 and peak-calling using MACS v2.1 (36). Experiments were repeated at least 3 times for each condition. Reads were counted within reproducible peaks and normalized to library size with DiffBind (37). Normalized read counts were log-transformed, and a linear batch effect between replicates was regressed out. Peaks with variable accessibility across groups were calculated using one-way ANOVA. We then applied k-means clustering (k=30) to the normalized, log-transformed, and batch-corrected read counts of variable peaks. Peaks that were insufficiently correlated with the average expression within each cluster (r < 0.7) were removed, and clusters with strongly correlated average expression (r > 0.85) were combined, resulting in 12 merged clusters. Motif enrichment analysis was performed with HOMER (11) using default parameters, comparing the peaks within each cluster to the background of all reproducible peaks in our dataset. Motifs were assigned to the TFs for which they are named, as well as to TFs with similar DNA-binding domains in the TFClass database, as previously described (38).

### Single-cell RNA-Sequencing

scRNA-Seq of the organoids was performed at D0 and D10 of differentiation to follow gene expression at the single cell level. Organoids were prepared from 6-12 newborn ears of both sexes, and more than 5,000 cells were collected for analysis using the 10X Genomics droplet-based single-cell sequencing platform. The cell suspension was diluted to a concentration of 500 cells per ml and immediately captured, lysed, and primed for reverse transcription (RT) using the high throughput, droplet microfluidics Gemcode platform from 10X Genomics with v2 chemistry. Each droplet on the Gemcode co-encapsulates a cell and a gel bead that is hybridized with oligo(dT) primers encoding a unique cell barcode and unique molecular identifiers (UMIs) in lysis buffer. After capture for 6 min on gel beads, the transcriptomes were pooled and reverse transcribed to cDNA. Cell barcodes and UMIs were employed, after sequencing to demultiplex the originating cell and mRNA transcript from the pooled and PCR amplified cDNA. RT-PCR amplification of cDNA, and preparation of a library from 30 ends were conducted according to the manufacturer’s published protocol. We performed 14 cycles of PCR amplification of cDNA. The library was sequenced on an Illumina NovaSeq 6000 with an S2 100 cycle reagent kit at the Broad Institute Sequencing Facility.

### Single cell RNA-seq pre-processing, integration, and clustering

Initial scRNA-seq data processing, including demultiplexing, alignment to the mouse genome (mm10), and read counting were performed with cellranger (10X Genomics). The number of genes expressed, the number of UMIs detected, and the percentage of mitochondrial and ribosomal RNA were calculated for quality control. Cells with >5% of UMIs from mitochondrial genes were discarded. We applied the Seurat v3 Standard Workflow (39) to integrate cells across replicates, using 7,000 highly variable genes, 3000 anchors, and 50 dimensions. Subsequently, principal component analysis was performed, and cells clustered on a K-nearest neighbor graph based on Euclidean distance using the previously defined PCA dimensionality as the input. Cells were clustered using the Louvain algorithm to optimize the standard modularity function before performing dimensionality reduction via UMAP. Further analysis was performed by re-clustering selected sets of cells followed by differential gene expression analysis to identify unique cell markers.

scRNA-seq for postnatal day 2 and day 7 mouse utricle was performed as follow: 3 mice (CD-1 background) were euthanized and their temporal bone removed. Utricles were harvested and incubated in thermolysin (Sigma-Aldrich) for 20 min at 37°C. Thermolysin was then replaced with Accutase (Sigma-Aldrich) and the tissue incubated for 3 min at 37° C followed by mechanical dissociation until a single cell suspension was obtained. After inactivation of the Accutase with 5% fetal bovine serum, the cell suspension was filter through a 35μm nylon mesh and processed for scRNA-seq at the Institute for Genome Sciences (IGS) of the University of Maryland, School of Medicine. Approximately 10,000 dissociated utricular cells were captured into a Chromium Controller (10x Genomics) for droplet-based molecular barcoding. Library preparation was performed using the 10x Single Cell Gene Expression Solution. Libraries from two utricular samples were sequenced across three lanes of an Illumina HiSeq4000 sequencer to produce paired-end 75 bp reads.

### Comparison of organoid cell clusters to in vivo cell types

Cell clusters in organoids were compared to cell types in the *in vivo* mouse cochlea based on correlations among shared marker genes. A cell type specificity score was defined for each gene in each cluster, as previously described (40). Briefly, for each gene detected in >50% of cells from a cluster, we multiplied its enrichment (log_2_ fold change) by its specificity (percent of relevant cluster expressing the gene/percent of other clusters expressing the gene) to yield a specificity score within each cluster. Specificity scores were calculated for each organoid cell cluster, as well as for each cell type in the mouse cochlea, based on scRNA-seq of E14, E16, P1, and P7 cochlear cell types (5). We then used Pearson’s correlations to quantify the similarity of marker genes for each *in vivo* vs. *in vitro* cell cluster.

In addition, we used the projectR R package (41) to “score” each cell for the expression of marker genes in each in vivo cell type. For each in vivo cochlear cell type, we defined a set of specifically expressed markers (>50% non-zero counts; p-value < 0.05). Cells in organoids were then scored based on the combined expression of each set of marker genes, using the projectR function. Similarly, cells in organoids were compared to other cell types that share an Atoh1 lineage, defining scores based on genes specifically expressed in gut cells, Merkel cells, cerebellar granule cell progenitor cells, and hair cells (6).

### Monocle trajectory construction

We reconstructed a pseudotime trajectory for the differentiation of organoids to hair cells using Monocle. For this analysis, we selected the organoid cluster most similar to greater epithelial ridge (cluster 4), as well as the two clusters of hair cells. Trajectory analysis was performed using Monocle version 2.14.0. Cells were ordered using the top 375 differentially expressed genes (min.pct = 0.25) in the cells from each cluster. Monocle’s orderCells function arranged these cells along a pseudotime trajectory. The differentialGeneTest function (fullModelFormulaStr = “sm.ns(Pseudotime)”) was used to calculate the significance of each gene’s expression change over pseudotime. Genes with a q-value < 0.01 and detected in at least 200 cells were retained for clustering in pseudotime.

### GO term enrichment

GO term enrichment on the genes supported by the bulk RNA-seq clusters and the pseudotime-derived scRNA-seq clusters was performed using clusterProfiler R package (41). GO terms with a BH adjusted p-value < 0.05 were reported.

### Gene regulatory network reconstruction

A gene regulatory network for the differentiation of organoids to hair cells was derived from scRNA-seq data. As with trajectory analysis, we selected clusters 4, 7, and 8 for this analysis. First, GENIE3 (9) was used to predict target genes for each of 1,186 transcription factors, the subset of transcription factors with expression in these cells from a list of 1675 mouse transcription factors from http://genome.gsc.riken.jp/TFdb/data/tf.name. The GENIE3 output was a list of transcription factors and their predicted target genes (regulons). We used hypergeometric tests to identify transcription factors whose predicted target genes were over-represented in each gene co-expression cluster from pseudo-time analysis and bulk RNA-seq. We further reconstructed a dynamical model for TF-to-TF interactions during hair cell differentiation using SCODE, which implements an ordinary differential equation model using pseudotime. For this analysis, we selected 58 TFs whose targets were over-represented within gene co-expression clusters whose expression peaked early, middle, or late in hair cell differentiation, consistently in our bulk and single-cell datasets.

### Data availability

The bulk RNA-seq, scRNA-seq, and ATAC-seq data from this study are available on GEO (accession number GSE132635) and in an interactive version on gEAR at umgear.org/Lgr5org (Supplemental Figure 3) (42).

## SUPPLEMENTAL INFORMATION

Supplemental Information includes three figures and eleven tables.

## AUTHOR CONTRIBUTIONS

G.K., D.L., D.A., C.H., R.H., S.A.A. and A.E. designed research; G.K., D.L., D.A. and C.H. performed research; G.K., D.L., D.A., C.H., B.R.H., C.C., B.M., M.S., A.C.S., R.H., R.A.S., S.A.A. and A.E. analyzed data; and G.K., D.L., D.A., R.H., R.A.S., S.A.A. and A.E. wrote the paper.

## ACKNOWLEDGEMENTS

We thank the Dana-Farber Genomics Core, the Harvard Chan Bioinformatics Core, and the Broad Institute Sequencing Facility for technical assistance and the NIH (grant DC007174) and the Hearing Restoration Project of the Hearing Health Foundation for funding.

## Supplemental Figures

**Supplemental Figure 1 (relevant to Figure 5).**
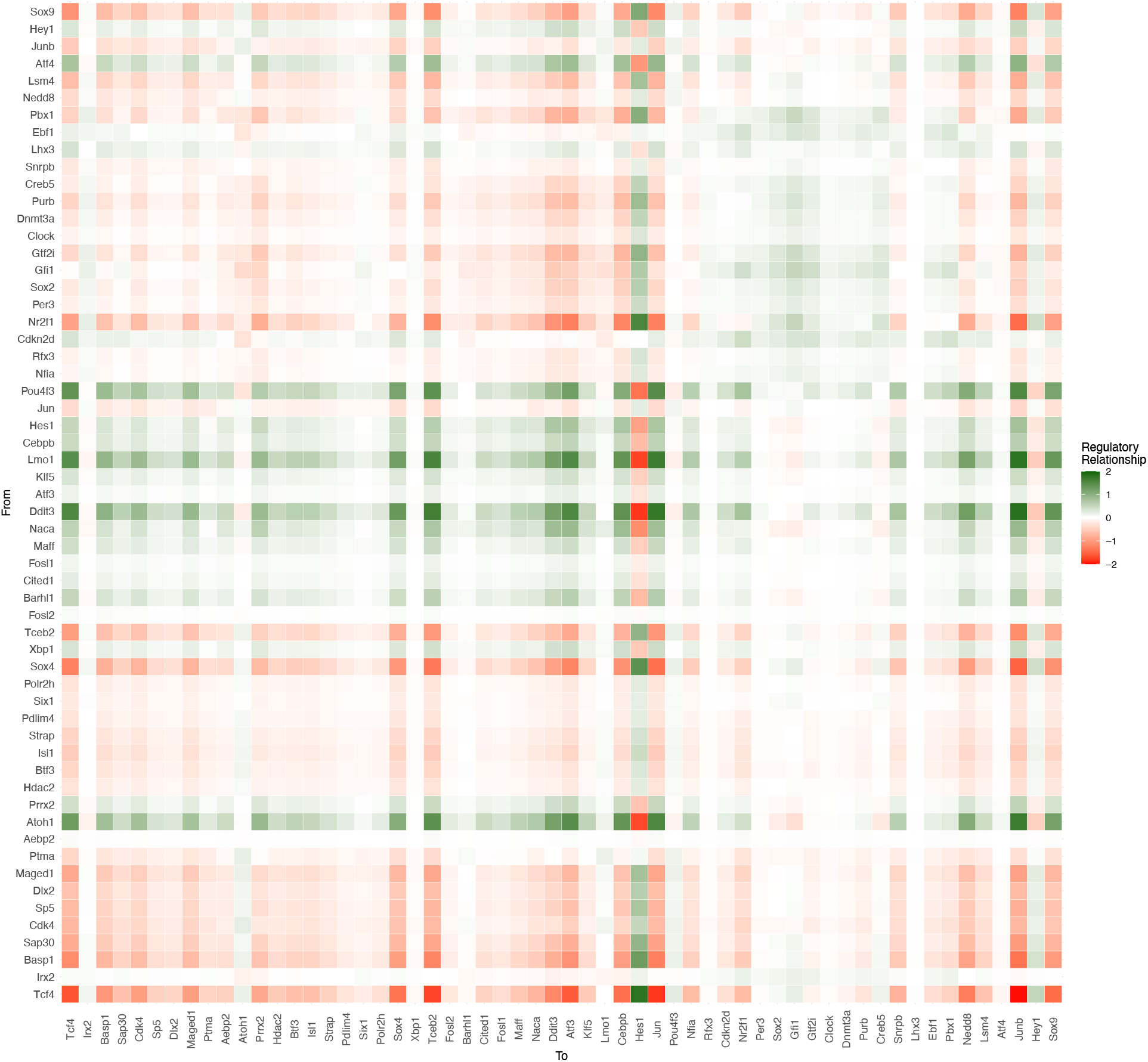
Heatmap of the highly connected genes in the gene regulatory network.

**Supplemental Figure 2 (relevant to Figure 6).**
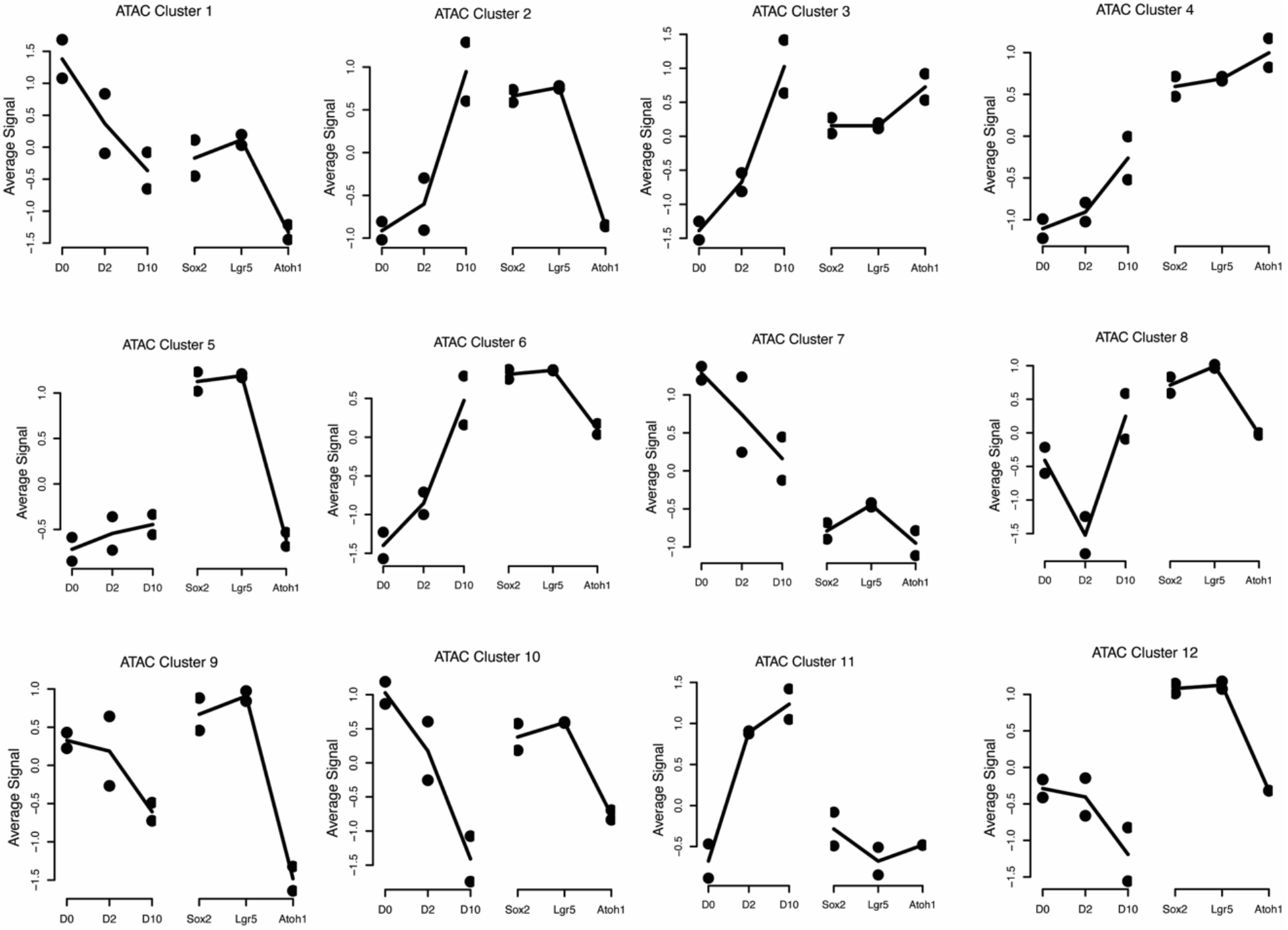
K-means clustering of patterns of chromatin opening from the ATAC-seq data of the differentiating cochlear organoids at D0, D2 and D10.

**Supplemental Figure 3 (relevant to Figures 2, 3 and 6).**
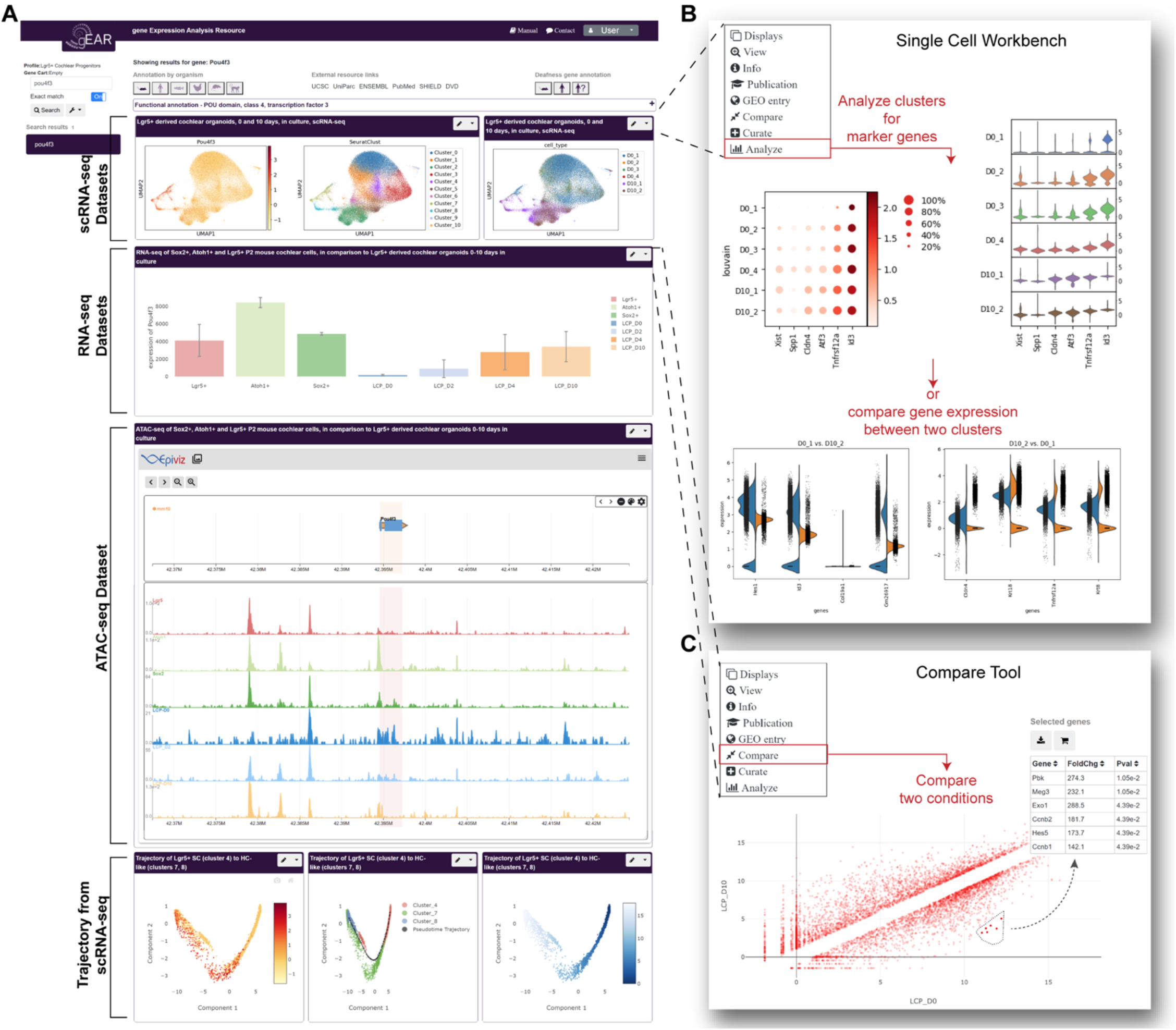
A-C. A custom profile was generated in the gEAR to support sharing, visualization and analysis of the processed data presented in this manuscript (https://umgear.org/Lgr5org). Overview (A)of the manuscript profile, which contains the scRNA-seq data (UMAP), the RNA-seq data (bar graph), the ATAC-seq data (Epiviz) and trajectory. Example of the single cell workbench (B) showing marker genes for the two time points across biological replicates (top) and the top 4 differentially expressed genes between day 0 replicate 1 (D0_1) and day 10 replicate 2 (D10_2) (bottom). Example of the use of the ‘compare tool’ (C), showing the differentially expressed genes between D0 and D10 time points.

